# Zelda and the evolution of insect metamorphosis

**DOI:** 10.1101/368035

**Authors:** Alba Ventos-Alfonso, Guillem Ylla, Xavier Belles

## Abstract

In the Endopterygote *Drosophila melanogaster*, Zelda is a key activator of the zygotic genome during the maternal-to-zygotic transition (MZT). Zelda binds *cis*-regulatory elements (TAGteam heptamers), and makes chromatin accessible for gene transcription. Recently, Zelda has been studied in two other Endopterygotes: *Apis mellifera* and *Tribolium castaneum*, and the Paraneopteran *Rhodnius prolixus*. We have studied Zelda in the cockroach *Blattella germanica*, a hemimetabolan, short germ-band, and Polyneopteran species. Zelda protein of *B. germanica* has the complete set of functional domains, which is typical of lower insects. The TAGteam heptamers of *D. melanogaster* have been found in the *B. germanica* genome, and the canonical one, CAGGTAG, is present at a similar relative number in the genome of these two species and in the genome of other insects, suggesting that, although within certain evolutionary constraints, the genome admits as many CAGGTAG motifs as its length allows. Zelda-depleted embryos of *B. germanica* show defects involving the blastoderm formation and the abdomen development and have genes contributing to these processes down-regulated. We conclude that in *B. germanica* Zelda strictly activates the zygotic genome, within the MZT, a role conserved in more derived Endopterygote insects. In *B. germanica, Zelda is expressed during MZT, whereas in D. melanogaster* and *T. castaneum* it is expressed well beyond this transition. Moreover, in these species and *A. mellifera*, Zelda has functions even in postembryonic development. The expansion of Zelda expression and functions beyond the MZT in holometabolan species might have been instrumental for the evolutionary transition from hemimetaboly to holometaboly. In particular, the expression of Zelda beyond the MZT during embryogenesis might have allowed building the morphologically divergent holometabolan larva.

**Author summary:** In early insect embryo development, the protein Zelda is a key activator of the zygotic genome during the maternal-to-zygotic transition. This has been thoroughly demonstrated in the fruit fly *Drosophila melanogaster*, as well as in the red flour beetle *Tribolium castaneum*, both species belonging to the most modified clade of endopterygote insects, showing complete (holometabolan) metamorphosis. In these species, Zelda is expressed and have functions in early embryogenesis, in late embryogenesis and in postembryonic stages. We have studied Zelda in the German cockroach, *Blattella germanica*, which belong to the less modified clade of polyneopteran insects, showing an incomplete (hemimetabolan) metamorphosis. In *B. germanica*, Zelda is significantly expressed in early embryogenesis, being a key activator of the zygotic genome during the maternal-to-zygotic transition, as in the fruit fly and the red flour beetle. Nevertheless, Zelda is not significantly expressed, and presumably has no functions, in late embryogenesis and in postembryonic stages of the cockroach. The data suggest that the ancestral function of Zelda in insects with hemimetabolan metamorphosis was to activate the zygotic genome, a function circumscribed to early embryogenesis. The expansion of Zelda expression and functions to late embryogenesis and postembryonic stages might have been a key step in the evolutionary transition from hemimetaboly to holometaboly. In hemimetabolan species embryogenesis produces a nymph displaying the essential adult body structure. In contrast, embryogenesis of holometabolan species produces a larva that is morphologically very divergent from the adult. Expression of Zelda in late embryogenesis might have been a key step in the evolution from hemimetaboly to holometaboly, since it would have allowed the building the morphologically divergent holometabolan larva.

## Introduction

An important event in early embryo development in metazoans is the maternal-to-zygotic transition (MZT), by which maternal mRNAs are eliminated and the zygotic genome becomes activated and governs further development [1]. In the MZT context, the protein Zelda was found to be a key activator of the early zygotic genome in the fruit fly, *Drosophila melanogaster* [2], an Endopterygote species that shows a long germ-band embryo development and an holometabolan mode of metamorphosis.

The history of Zelda, however, starts in 2006, when Staudt et al. [3] reported a X-chromosomal gene in *D. melanogaster* encoding a nuclear zinc-finger protein, whose transcripts were maternally loaded and ubiquitously distributed in eggs and preblastoderm embryos. Loss of gene activity disrupted the pattern of mitotic waves in preblastoderm embryos, elicited asynchronous DNA replication, and caused improper chromosome segregation during mitosis. The gene was named *vielfältig*, the German word for versatile or manifold [3]. The same year, ten Bosch et al. [4] described that the heptamer CAGGTAG (a consensus sequence of a larger series of similar heptamers collectively referred to as TAGteam) is overrepresented in regulatory regions of many of the early transcribed genes in the zygote of *D. melanogaster*. Subsequently, Liang et al. [2] described the protein that binds specifically to the TAGteam sites and can activate transcription in transfection assays. The protein turned out to be Vielfältig, but the authors renamed it as Zelda (from <u>Z</u>inc-finger <u>e</u>arly <u>D</u>rosophila <u>a</u>ctivator). *D. melanogaster* embryos lacking Zelda showed defects in the formation of the blastoderm and could not activate genes essential for cellularization, sex determination, and pattern formation. Liang et al. [2] concluded that Zelda played a key role in the activation of the early zygotic genome of D. melanogaster.

Three years later, two back-to-back papers showed that during early embryogenesis of *D*. *melanogaster* Zelda marks regions subsequently activated during the MZT and promotes transcriptional activation of early-gene networks by binding to more than 1,000 DNA regulatory regions [5,6]. It became soon clear that the most important role of Zelda in *D. melanogaster*, and from which many of the observed effects derive, is to increase the accessibility of chromatin, allowing the transcription of multiple genes [7–9]. Associated to these observations, it was shown that around the MZT, Zelda is required for the appearance of topologically associated domain boundaries, which influence chromatin architecture and activation of gene expression [10].

From the point of view of the functional organization, a first work by Hamm et al. [11] showed that the protein Zelda of *D. melanogaster* presents a low complexity region, corresponding to the transactivation domain, and four C-terminal Zinc fingers, which are required for DNA binding and transcriptional activation [11]. More recently, Ribeiro et al. [12] reported that Zelda is conserved throughout insects and crustaceans and possess other conserved regions, including an acidic patch (which could be involved in the recruitment of chromatin remodeling proteins) and two N-terminal Zinc fingers [12]. Subsequently, Hamm et al. [13] approached the functional study of Zelda conserved domains using a Cas9 engineering system to obtain *D. melanogaster* insects with point mutations in the Zinc finger 1, in the acidic patch, or in the Zinc finger 2 (JAZ). Among other results, the experiments showed that mutations in the Zinc finger 1 or in the acidic patch did not affect the viability of the insects, whereas mutations in the Zinc finger 2 (JAZ) led to a maternal-effect lethal phenotype [13]. An additional transcriptomic analysis of mutated flies showed that the Zinc finger 2 (JAZ) acts as a maternal repressive domain during embryo development, being associated with the transcriptional regulation of the zygotic Zelda targets and with the clearance of maternal transcripts [13].

Outside *D. melanogaster*, expression of Zelda has been studied in the honeybee *Apis mellifera*, another Endopterygote, long germ-band, holometabolan species. Expression was observed in early embryogenesis in relation to blastoderm formation and gastrulation [14]. These expression features and the occurrence of TAGteam heptamers in *A. mellifera*, suggest that in this species Zelda plays the same role of activator of the early zygotic genome described in *D. melanogaster*. Moreover, Pires et al. (2016) also found Zelda expression in late embryogenesis of *A. mellifera*, associated to the central nervous system (CNS) precursor cells. This suggests that in the honeybee, Zelda contributes to the formation of the CNS, as occurs in *D. melanogaster* [15]. Also outside *D. melanogaster*, a recent contribution of Ribeiro et al. [12] added new information about the organization of the protein, showing that Zelda was an innovation of the Pancrustacea lineage. Beyond that, the study affords functional information on an Endopterygote and holometabolan species but that shows short germ-band embryo development, the red flour beetle *Tribolium castaneum*. Using RNAi approaches, Ribeiro et al. [12] demonstrate that Zelda is key not only during the MZT, but also in posterior segmentation and patterning of imaginal disc structures of *T. castaneum*. Additionally, they also show that Zelda is important for posterior segmentation in the kissing bug *Rhodnius prolixus* [12], a Paraneopteran, short germ-band and hemimetabolan species.

In a recent work where we compared broad transcriptomic differences along the ontogeny of *D. melanogaster* and the German cockroach, *Blattella germanica (a poorly* modified Polyneopteran, hemimetabolan and short germ-band species), Zelda emerged as a differential factor [16]. In *B. germanica*, the pattern of expression showed to be sharply concentrated in early embryo development, whereas in *D. melanogaster* Zelda expression covered the entire embryogenesis [16]. We related this difference with the modes of metamorphosis of the studied species, hemimetabolan in *B. germanica* and holometabolan in *D. melanogaster*. Hemimetaboly is characterized by an embryogenesis that produces a nymph displaying the essential adult body structure. The nymph grows through successive molts, and during the final molt to the adult stage the formation of functional wings and genitalia is completed. In holometabolan species, embryogenesis gives rise to a larva with a body structure considerably divergent from that of the adult, often more or less vermiform. The larva grows through various stages until molting to the pupal stage, which bridges the gap between the divergent larval morphology and that of the winged and reproductively competent adult [17]. We speculated that the difference in Zelda expression pattern would have to do with the different modes of metamorphosis. Thus, transient expression in very early embryo of *B. germanica* would be the ancestral condition in hemimetabolan species, which would trigger a discrete activation of the zygotic genome; conversely, continuous expression of Zelda during the embryogenesis of *D. melanogaster* might be needed for morphogenesis of the derived body form of the holometabolan larva [16]. The present work aims at illuminating these speculations by examining in *B. germanica* the organization of the protein, the distribution of TAGteams in the genome, the expression of Zelda along the ontogeny, and their functions, explained in morphological and molecular terms.

## Results

### Zelda organization is conserved in *Blattella germanica*

By combining BLAST search in *B. germanica* transcriptomes, mapping the resulting sequences in the *B. germanica* genome, and PCR strategies, we obtained a cDNA of 2,754 nucleotides comprising the complete ORF (GenBank accession number LT71728.1), which once translated gave a 918 amino acid sequence with high similarity to Zelda proteins, which we called BgZelda. Analyzing our 11stages RNA-seq dataset [16], we concluded that BgZelda gene do not show splicing isoforms involving the coding region, thus it produces a single transcript, which contrasts with the different amino acid sequences described for the Zelda gene of *D. melanogaster [11,18]*.

The canonical Zelda gene originally reported in *D. melanogaster*, DmZelda, is described as composed by a JAZ-C2H2 zinc finger domain and four C2H2 zinc fingers in the C-terminus region [2,11,18]. However, a recent work on insect and crustacean Zelda sequences [12] reported two additional zinc finger domains upstream the Zinc finger 2 (JAZ), although some insects have lost the first of them. Following the nomenclature of Ribeiro et al (2017), BgZelda possess all seven C2H2 zinc fingers: two of them at the N-terminal part of the protein (ZF-Novel and ZF1), then the ZF2 (JAZ domain) follows, and at the C-terminal region there are the other 4 ZFs (ZF3 to ZF6). Between ZF-Novel and ZF1 there is the “patch” region, with the sequence TMAPADSSD, and the “conserved region” with the typical motif RYHPY. Then, between ZF1 and ZF2 we found the “acidic patch”, characterized by the motif [DE]I[LW]DLD, which in the case of BgZelda is EILDLD. Finally, between ZF2 and ZF3, there is the “conserved region”, which in BgZelda has the sequence PPNLMAGPPISMEAQTEGLP, and the “activation region”, with the motif LPSFAQ (Fig 1A). A high conservation was found in the C-terminal four zinc fingers ZF3 to ZF6, which are the responsible for recognizing and binding to the DNA TAGteam heptamers, as demonstrated in *D. melanogaster* [11]. In this ZF3 to ZF6 region the percentage of similarity between DmZelda and BgZelda is 76.7% (S1 Fig).

**Fig 1.**
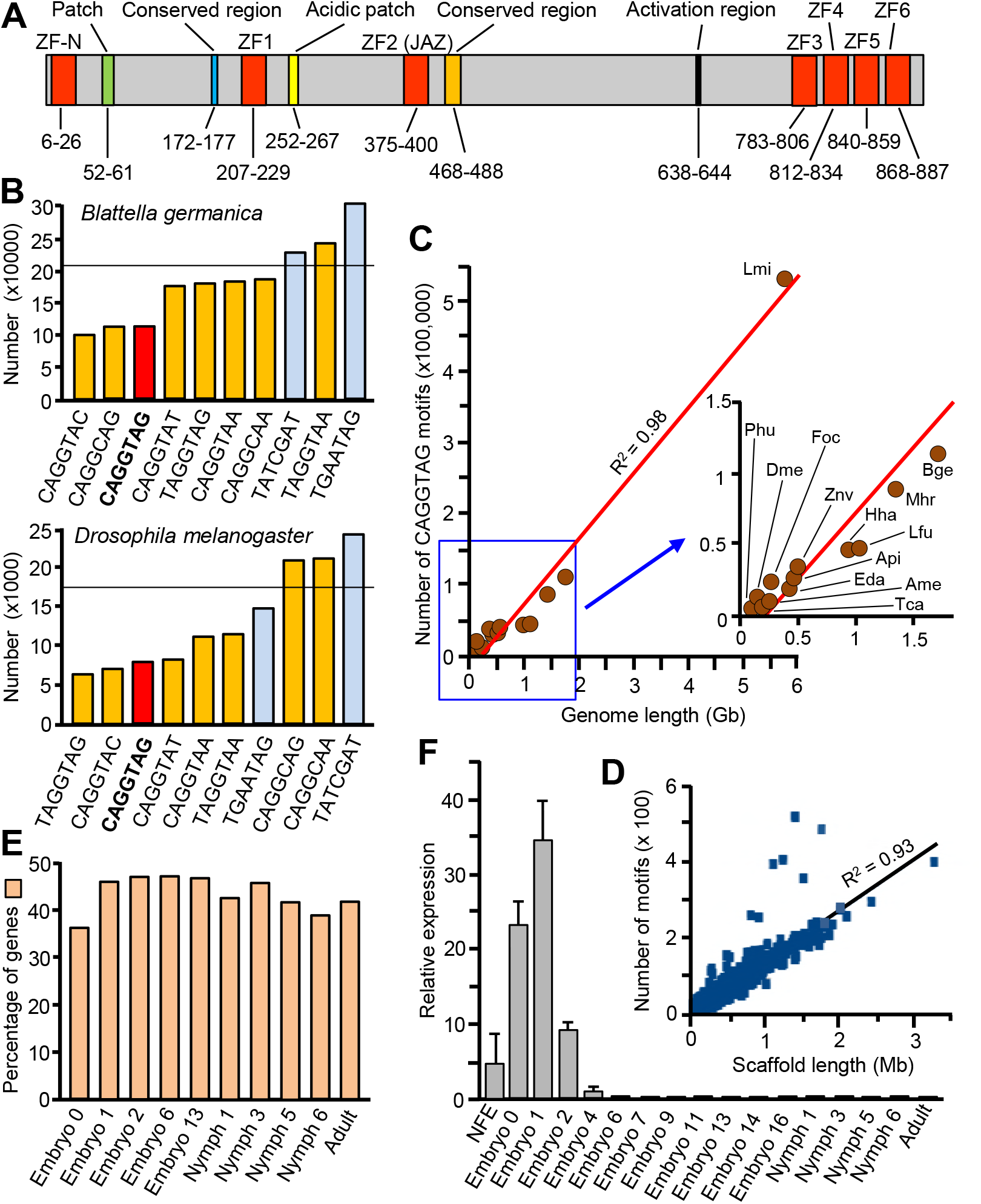
Structure and expression of Zelda in *Blattella germanica*, and the TAGteam heptamers. (A) Structure of the protein Zelda showing the most characteristic motifs according the nomenclature of Ribeiro et al. (2017); the numbers indicate the initial and final starting amino acid of each motif. (B) Occurrence of the eight functional (yellow and red columns, the red column indicates the canonical sequence CAGGTAG) and not functional (blue columns) TAGteam heptamers reported by Nien el al (2011) in *D. melanogaster* and *B. germanica* genomes; the horizontal line crossing the columns indicates the expected occurrences of a random heptamer. (C) Correlation between genome length and number of CAGGTAG motifs in hexapods; the genomes included are the Archaeognatha *Machilis hrabei* (Mhr), the Odonata *Ladona fulva* (Lfu), the Ephemeroptera *Ephemera danica* (Eda), the Orthoptera *Locusta migratori*a (Lmi), the Blattodea *B. germanica* (Bger), the Isoptera *Zootermopsis nevadensis* (Znv), the Hemiptera *Halyomorpha halys* (Hha), *Acyrthosiphon pisum* (Api) and *Frankliniella occidentalis (Foc), the Phthiraptera Pediculus humanus* (Phu), the Hymenoptera *Apis mellifera* (Ame), the Coleoptera *Tribolium castaneum* (Tca), and the Diptera *D. melanogaster* (Dme); the shown regression line has a Pearson correlation coefficient of 0.98. (D) Length of the scaffolds of the *B. germanica* genome (masked regions excluded) and number of CAGGTAG motifs in each scaffold; the shown regression line has a Pearson correlation coefficient of 0.93. (E) Fraction of genes containing the CAGGTAG heptamer among those up-regulated at each stage in the 11 stage RNA-seq libraries reported by Ylla et al. (2018) that cover the entire ontogeny of *B. germanica*; the genes up-regulated in “Embryo Day 0” were computed using the previous “Non-fertilized eggs” (NFE) library as a reference; superimposed is shown the pattern of expression of Zelda obtained from these libraries (see Ylla et al., 2018, for more details on the libraries). (F) Expression pattern of Zelda along ontogeny in *B. germanica* obtained by qRT-PCR; the stages examined are the same of the libraries of Ylla et al (2018), plus a number of additional embryo stages (ED4, ED7, ED9, ED11, ED14, ED16); each value represents 3 biological replicates and it is represented as copies of Zelda mRNA per 100 copies of BgActin-5c mRNA (mean ± SEM).

### *D. melanogaster* TAGteam heptamers are present in *Blattella germanica* genome

In *D. melanogaster, the C-terminal cluster of four zinc fingers recognizes particular cis-* regulatory heptamer motifs collectively known as TAGteam, being CAGGTAG the one to which Zelda bound with the highest affinity [2,4,5]. Nien et al. (2011) experimentally assessed that DmZelda binds the sequence CAGGTAG and, to a lesser extent, the seven additional TAGteam heptamers: TAGGTAG, CAGGTAC, CAGGTAT, CAGGTAA, TAGGTAA, CAGGCAG and CAGGCAA. Conversely, two additional heptamers (TGAATAG and TATCGAT) were not bound by DmZelda. In the *D. melanogaster* genome we found 95,495 heptamers that coincided with any of the eight DmZelda-binding TAGteam elements identified by Nien et al. (2011), whereas 1,299,345 were found in the longer genome of *B. germanica*. Significantly, these amounts are clearly lower than those expected if the occurrence of these eight heptamers were by chance (68 and 78% of the expected by chance, respectively). The canonical heptamer CAGGTAG appears 8,068 times in the *D. melanogaster* genome and 112,830 in that of *B. germanica* (Fig 1B) (45 and 50% of the expected by chance, respectively). Interestingly, the number of CAGGTAG heptamers appears roughly proportional to the length of the genome (1 per 15,160 nt in *B. germanica* and 1 per 17,894 nt in *D. melanogaster*). To ascertain whether this proportionality is more general in insects, we searched the number of CAGGTAG heptamers in the genome of thirteen insect species from different orders, with genome lengths ranging from 110 Mb (*Pediculus humanus*) to 5.7 Gb (*Locusta migratoria*) (S1 Table). The analysis revealed a strong and significant correlation (R^2^=0.98 and Pearson correlation coefficient, P-value<0.05) between genome length and number of CAGGTAG heptamers (Fig 1C). Moreover, we found that the number of CAGGTAG heptamers in each scaffold of the *B. germanica* genome is proportional to the length of the scaffold (Fig 1D and S2 Table) and are distributed more or less regularly within the scaffold (S2 Table). This suggests that the 112,830 CAGGTAG motifs found in *B. germanica* are distributed more or less regularly throughout the genome, without significant concentrations in given regions.

Finally, we wanted to examine the pattern of expression of the genes containing the CAGGTAG heptamer in the promoter region. For this purpose, we identified the *B. germanica* genes having at least one CAGGTAG motif within the 2 kb upstream the start codon, a stretch than we considered that would contain the promoter region or a major part of it. Then, using the 11 stage RNA-seq libraries that cover the entire ontogeny of *B. germanica* [16] we obtained the fraction of genes containing the CAGGTAG motif in the promoter region among those up-regulated at each stage. Results showed that this gene fraction is quite constant during the successive developmental transitions (Fig 1E).

### BgZelda is expressed in a narrow temporal window around the MZT

Using qRT-PCR, we studied the expression of BgZelda along the ontogeny of *B. germanica* at the 11 stages used in the RNA-seq libraries reported by Ylla et al. [16], plus five additional embryo stages (days 7, 9, 11, 14 and 16) in order to cover the embryogenesis more thoroughly. Results show that BgZelda is predominantly expressed in the first stages of embryogenesis, between day 0 and day 2, with a clear peak on day 1 (35 copies of BgZelda as average per 100 copies of actin), that is within the MZT. Beyond embryo day 6 and until the adult stage, BgZelda is expressed at comparatively very low levels (between 0.006 and 0.08 copies of BgZelda as average per 100 copies of actin) (Fig 1F).

Significantly, the pattern obtained using qRT-PCR measurements is very similar to that obtained by computing the reads (FPKM) from the 11 stages of the RNA-seq libraries [16] (S3 Fig). Indeed, a remarkable correlation (R^2^= 0.82) is obtained when using the same 11 stage points and comparing the FPKM (S3 Fig) and the qRT-PCR (Fig 1E) values. The expression of BgZelda concentrated at the beginning of embryogenesis (Fig 1F and S3 Fig), contrasts with the expression in *D. melanogaster* obtained from transcriptomic data [16], which spans all along embryogenesis and first larval instar and then declines during the remaining post-embryonic stages (S3 Fig). Northern blot analyses had shown that DmZelda expression appears quite high in the embryo, the first two larval instars, then decreases in the third larval instar and the pupa, and slightly increases in the adult [3], which is fairly coincident with the FPKM pattern obtained by us (S3 Fig). The analysis of transcriptomic data of *T. castaneum* that covers the first 30% of the embryogenesis divided into four successive equal intervals [19], suggests that TcZelda is also expressed in a wide time frame. Data shows that expression of TcZelda progressively increases at least until 48h after egg laying (corresponding to 30% embryo development), which is more similar to the pattern of *D. melanogaster* (with a pattern of sustained expression during all embryogenesis) than that of *B. germanica* (with a sharp peak of Zelda expression at 6% embryo development) (S3 Fig).

### Depletion of BgZelda results in a diversity of embryo defects

Our functional study of BgZelda was approached through maternal RNAi. One-day-old adult females (AdD1) were injected with 3 µg of dsZelda (treated females), whereas equivalent females were injected with 3 µg of dsMock (control females). All females were allowed to mate and formed a first ootheca on AdD8. The oothecae of control females (n=32) gave normal embryos that hatched 19 days later (AdD27), producing in all 1,281 first instar nymphs (35–40 nymphs per ootheca). All the dsZelda-treated females (n=39), also produced a first ootheca on AdD8. Nineteen of the 39 oothecae (49%) gave normally hatched nymphs (35–40 nymphs per ootheca) in AdD27, whereas from the remaining 20 oothecae (51%) no nymphs hatched, or hatched very few (a maximum of 20 hatched nymphs per ootheca, but often much less). In total, from the 39 oothecae formed by dsZelda-treated females, we recorded 1024 nymphs that hatched normally and 210 embryos or nymphs that did not hatch, and that were dissected out from the totally or partially unviable oothecae on AdD28. These 210 embryos or unhatched nymphs showed wide range of phenotypes that were classified into the following seven categories in different stages defined according to Tanaka [20]. Phenotype A (Fig 2A): morphology of normal nymph in E19, thus ready to hatch, but which did not hatch. Phenotype B (Fig 2B): embryos in stage 16–18, morphologically similar to controls but showing and internal brown coloration. Phenotype C (Fig 2C): embryos in stage 17 but with the abdomen smaller than in controls. Phenotype D (Fig 2D): embryos in stage 16, but with the abdomen more elongated and smaller than in controls; this phenotype show the same essential defect as phenotype C, but being in an early developmental stage. Phenotype E (Fig 2E): embryos with the curvature of the body inverted, with the dorsal part concave instead of convex; they can reach the stage 15 and even beyond. Phenotype F (Fig 2F): embryos in medium stages of development, between stages 9 and 11, almost transparent and with the abdomen shorter than normal. Phenotype G (Fig 2G): embryos corresponding to very early development with no signs of segmentation. The most common (29% of abnormal embryos) was phenotype A, followed by D (23%), and then B, C, E and G ca. 10% penetrance each) and F (5%) (Fig 2H). If we merge categories C and D, as they show the same essential phenotype (small abdomen), then we can conclude that ca. 33% of the embryos from unviable oothecae present this defect.

**Fig 2.**
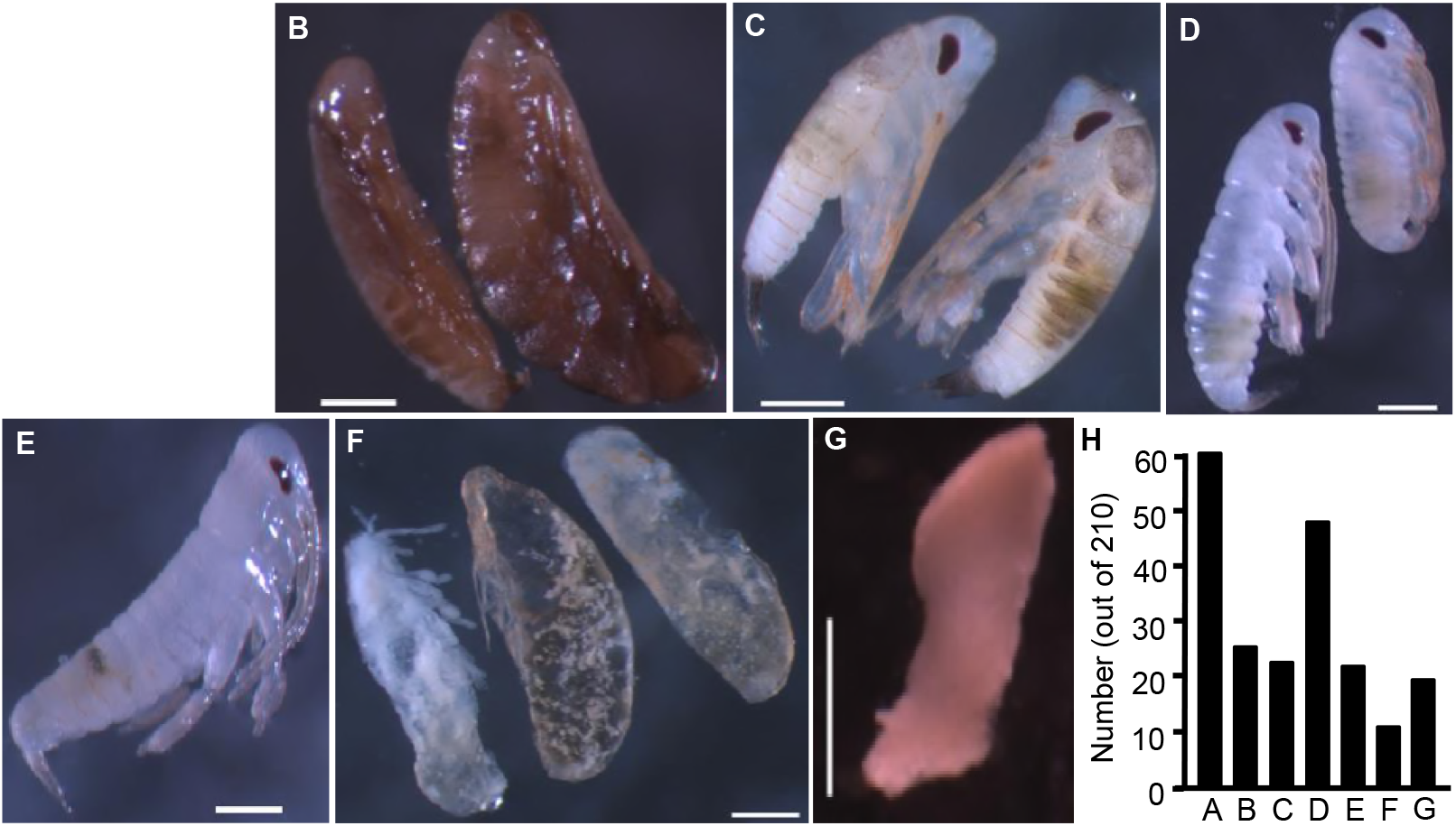
Different phenotypes observed in unhatched oothecae from Zelda-depleted *Blattella germanica*. (A) Phenotype A, which shows the morphology of an apparently normal first instar nymph ready to hatch, but which did not hatch. (B) Phenotype B. (C) Phenotype C. (D) Phenotype D. (E) Phenotype E. (F) Phenotype F. (G) Phenotype G. (H) Number of embryos showing the described phenotypes from a sample of 210 embryos studied. Scale bars in figures A-G: 500 µm.

As the most conspicuous expression of BgZelda extends from ED0 to ED2, and in ED4 already falls to very low levels (Fig 1F), we considered that BgZelda functions concentrates over that period. Consequently, we decided to study the BgZelda-depleted embryos just after the period of maximal expression, following the same protocol of maternal RNAi but dissecting the oothecae 4 days after its formation (at ED4 stage). Control females (n = 7) were treated with dsMock, and 172 ED4 embryos were studied from their oothecae, whereas from the oothecae of the dsZelda-treated females (n = 15) we examined 239 embryos. The 172 embryos from control females showed the normal aspect of an ED4 embryo (18–20% embryo development, Tanaka stage 4–5). From the 239 embryos examined in oothecae from dsZelda-treated females, a total of 99 (41%) were normal ED4 embryos, identical to controls (Fig 3A). The remaining 140 embryos showed a diversity of developmental delays and malformations, which were classified into 5 categories, as follows. Phenotype H (Fig 3B): Embryo with seriously impaired segmentation: the cephalic segments are more or less delimited, the three thoracic segments can be distinguished by three undulated furrows in the thorax region, and the abdominal region is amorphous. In terms of general shape, it resembles the Tanaka stages 2–3 (12–15% development). Phenotype I (Fig 3C): Embryo with seriously impaired segmentation, twisted, shorter and wider, and with the caudal region (which would correspond to the growth zone) amorphous and swollen. Phenotype J (Fig 3D): Embryo with no traces of segmentation, only furrows and undulations are insinuated in the cephalic and thoracic region. In terms of general shape, it resembles the Tanaka stages 1–2 (8–12% development). Phenotype K (Fig 3E): Embryo at the stage of germ band anlage, resembling the Tanaka stage 1 (4–8% development). Phenotype L (Fig 3F): Embryos developed in dorsal side of the egg instead of the ventral side. They show the shape of a Tanaka stage 5 (20% development), but with only the 3 or 4 first abdominal segments recognizable, the caudal part being amorphous. The most frequent (41% of abnormal embryos) was phenotype H, followed by J and K (ca. 20% each), and then I and L (ca. between 8 and 11%) (Fig 3G). Interestingly, the defect of abdomen malformation is common to all phenotypes, even in the less severe. Taken together, these results suggest that BgZelda depletion affects the elongation, segmentation and the formation of the abdomen, and the curvature of the body, possibly resulting from defects in the dorso-ventral (DV) patterning.

**Fig 3.**
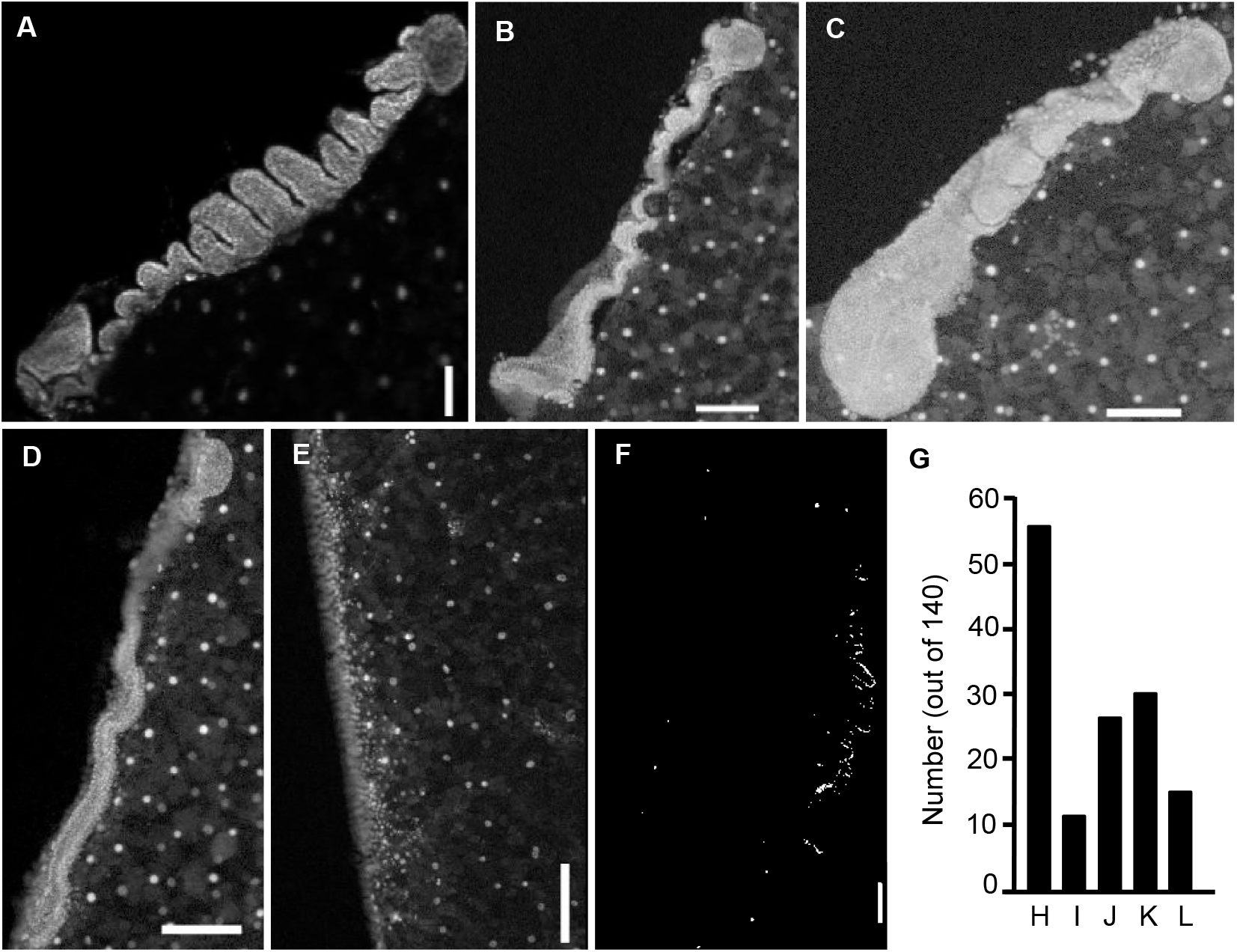
Different phenotypes observed in 4-day-old oothecae (embryos in stage ED2) from Zelda-depleted *Blattella germanica*. (A) Normal ED4 embryo. (B) Phenotype H. (C) Phenotype I. (D) Phenotype J. (E) Phenotype K. (F) Phenotype L. (G) Number of embryos showing the described phenotypes from a sample of 140 embryos studied. Scale bars in figures A-F: 200 µm.

Finally, although Zelda expression is very low in postembryonic stages (Fig 1F), we aimed at assessing if it can play some role in these stages. We focused on the metamorphic transition between N5 to N6 and to adult as the most relevant from a developmental point of view. Thus, freshly emerged fifth instar female nymphs (N5D0) were injected with 3 µg of dsZelda (treated nymphs, n = 15), whereas equivalent nymphs were injected with 3 µg of dsMock (controls, n= 12). All treated and control nymphs molted to morphologically normal N6, and subsequently to morphologically normal adults. The only difference between treated and controls was a slightly longer duration of the stages in the treated group. That of N5 was 6.0 + 0 days in controls and 6.13 + 0.35 days in treated, whereas that of N6 was 8.0 + 0 days in controls and 8.20 + 0.45 days in treated.

### Depletion of BgZelda impairs the expression of genes involved in early embryo development

The window of maximal expression of BgZelda is between ED0 and ED4, with an acute peak on ED1 (Fig 1E), and the most characteristic phenotypes of BgZelda-depleted specimens were observed in early embryogenesis (Fig 3). Therefore, we examined the expression of a number of genes involved in early embryo, at the ED2 stage, just after the peak of BgZelda expression. We started by assessing that the maternal RNAi was efficient, was measuring the expression of BgZelda in ED1 and ED2 oothecae from females treated with dsZelda or dsMock. Results showed that treatment with dsZelda reduced the BgZelda mRNA levels by 85% on ED1 and kept even lower levels in ED2 (Fig 4A).

**Fig 4.**
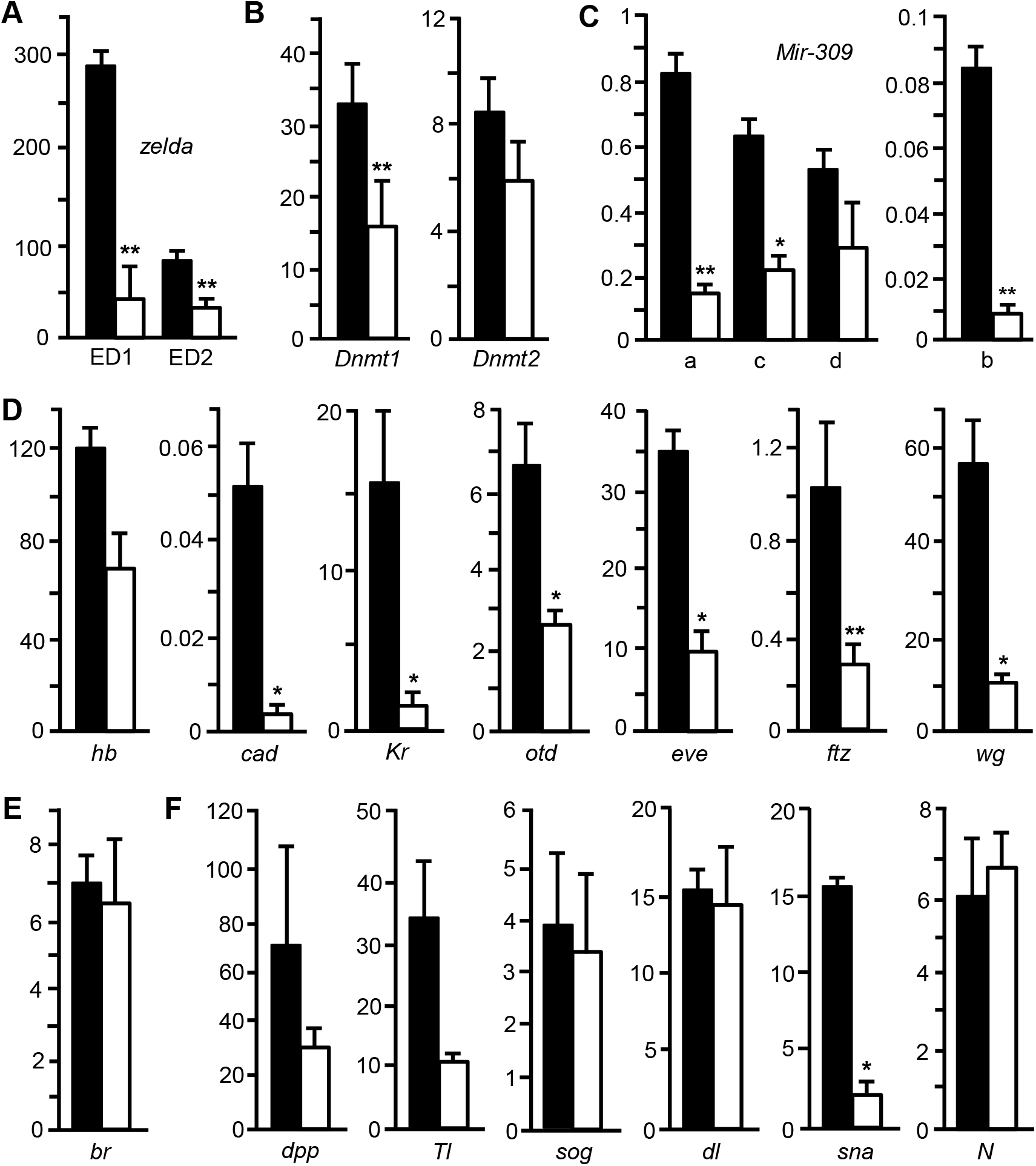
Effects of Zelda depletion on the expression of early embryogenesis genes in *Blattella germanica*. (A) Transcript decrease resulting from maternal RNAi of BgZelda, measured on ED1 and ED2. (B) Expression of *DNA-methyltransferase-1* (*DNMT1*) and *DNA-methyltransferase-2* (*DNMT2*). (C) Expression of the of the MIR-309 precursors. (D) Expression of the early patterning genes *hunchback* (*hb*), *caudal* (*cad*), *Krüppel* (*Kr*), *orthodenticle (otd*), *even-skipped* (*eve*), *fushi tarazu* (*ftz*) and *wingless* (*wg*) (E). Expression of *Broad complex* (*br*). (F) Expression of the genes related to dorso-ventral patterning *decapentaplegic* (*dpp*), *Toll* (*Tl*), *short-gastrulation* (*sog*), *dorsal* (*dl*), *snail* (*sna*) and *Notch* (*N*). The white column represents controls and the black column Zelda-depleted embryos. Each value represents 3 biological replicates and is expressed as copies of mRNA per 1000 copies of BgActin-5c mRNA (mean ± SEM). Asterisks indicate statistically significant differences with respect to controls (* p<0.05; **p<0.0005) calculated on the basis of Pair Wise Fixed Reallocation Randomization Test implemented in REST [37].

Subsequently, we examined the expression of a number of genes on ED2 in BgZelda-depleted and control embryos. In relation to epigenetic mechanisms, we measured the expression of *DNA-methyltransferase-1* (*Dnmt1*), involved in DNA methylation, and *DNA-methyltransferase-2* (*Dnmt2*), involved in tRNA methylation [21], and results indicated that *dnmt1* was significantly down-regulated whereas *dnmt2* was not (Fig 4B). Regarding microRNAs, it has been shown in *D. melanogaster* that DmZelda activates the expression of MIR-309 microRNAs in the MTZ context, which in turn eliminate maternal mRNAs [2,22]. We, thus, measured the expression of the precursors of the four variants of MIR-309 reported in *B. germanica* [23]. Results showed that, in all cases, the expression decreased in the BgZelda-depleted embryos at ED2 (Fig 4C). Then, given that a significant number of phenotypes showed defects in very early embryogenesis and in the formation of the abdomen, we studied a number of genes involved in these processes [24–27]. These included two maternal genes that are also expressed at the onset of zygotic activation: *hunchback* (*hb*) and *caudal* (*cd*), two gap genes: *Krüppel* (Kr) and *orthodenticle* (*otd*), two pair-rule genes: *even-skipped* (*eve*) and *fushi tarazu* (*ftz*), and one segment polarity gene: *wingless* (*wg*). Results indicated that the expression of most of these genes was significantly reduced in BgZelda-depleted embryos (Fig 4D). Moreover, previous studies have shown that maternal RNAi of *Broad Complex* (*br*) provoked embryo defects similar to those of BgZelda depletion, in particular deficiently developed abdomens [28]. Therefore, we also included *br* among the genes to be studied. However, results showed that BgZelda depletion did not affect *br* mRNA levels (Fig 4E). Although less frequent, another defect resulting from BgZelda depletion was the positioning of the embryo in the dorsal part of the egg. Thus, we studied the expression of a number of genes involved in dorso-ventral (DV) patterning, like *decapentaplegic* (*dpp*), *Toll* (*Tl*), *short-gastrulation* (*sog*), *dorsal* (*dl*), *snail* (*sna*) and *Notch* (*N*) [2,29]. Results showed that only the expression of *sna* resulted significantly reduced in BgZelda-depleted embryos (Fig 4F).

Additionally, we examined whether the genes whose expression was measured contained the consensus Zelda motif CAGGTAG, or any of the other seven TAGteam heptamers to which DmZelda binds: TAGGTAG, CAGGTAT, CAGGCAA, CAGGCAG, CAGGTAA, CAGGTAC and TAGGTAA [6]. Results indicate that *br, cad, dl, Dnmt2, hb, Kr, otd, Tl and N* have at least one CAGGTAG motif within the gene, and that all genes have at least one of the other seven TAGteam heptamers, except *cad* which has none (S3 Table). There is no apparent correlation between the occurrence of TAGteam motifs in the gene, and the effect of BgZelda on its expression. For example, *cad* has no TAGteam motifs and its expression appears stimulated by BgZelda, whereas *dl* has 32 motifs but BgZelda do not influence its expression (Fig 4 and S3 Table).

## Discussion

BgZelda has all the characteristic domains described by Ribeiro et al. (2017), including the ZF-Novel that was left in doubt for *B. germanica* in that work (Fig 1A). Ribeiro et al. (2017) found the same complete set of motifs in the Zelda ortholog of an Archaeognathan (*Machilis hrabei*), two Palaeopterans, (the Odonatan *Ladona fulva* and the Ephemeropteran *Ephemera danica*), and a basal Neopteran (the Isopteran *Zootermopsis nevadensis*). Then, one or more motifs are absent in different species of higher clades, whereas the set is very incomplete in the Zelda orthologs of non-insect hexapods and in crustaceans [12]. This suggests that the complete set of Zelda motifs is an ancestral condition in insects, which is still present in Archaeognatha, Palaeoptera and basal Neoptera like *B. germanica*, while some of the motifs became secondarily lost in different lineages of Paraneoptera and Endopterygota.

In the genome of *B. germanica*, we have been found all the TAGteam heptamers to which DmZelda binds in *D. melanogaster*. Interestingly, the canonical one, CAGGTAG, is present at a similar relative number in these two and in other insect genomes studied genomes (Fig C). This suggests that, although within certain evolutionary constraints, the genome admits as many CAGGTAG motifs as its length allows. Moreover, the distribution along the genome, at least in *B. germanica*, is very regular, without accumulations or biases in given regions, in general (Fig 1D, S2 Fig and S2 Table). The whole data suggest that in *B. germanica*, Zelda also pays the role of pioneer factor that binds DNA regions through these motifs, making the chromatin accessible for transcription, which had been postulated for Endopterygotes like *D. melanogaster* [5,30].

BgZelda is mostly expressed in a narrow window in early embryogenesis, with an acute expression peak on ED1, within the MZT (Fig 1F). This contrasts with the pattern observed in *D. melanogaster and T. castaneum*, where high expression is maintained beyond the MZT (S3 Fig). These two latter species are holometabolan, but embryo develops through short germ-band mode in *T. castaneum* and long germ-band in *D. melanogaster*. Thus the different pattern of expression of Zelda in *B. germanica* with respect to *D. melanogaster* and *T. castaneum* appear to relate with the metamorphosis mode. Hemimetabolan species, like *B. germanica*, develop the basic adult body structure during embryogenesis, whereas holometabolan species, like *D. melanogaster* and *T. castaneum*, delay adult body building until the pupal stage [17,31]. Thus, the expression of Zelda beyond the MZT in *D. melanogaster and T. castaneum*, might be needed to activate successive gene sets needed to build, during embryogenesis, the divergent holometabolan larval morphology. Moreover, Zelda expression and functions significantly extend beyond the embryo in holometabolan species (S3 Fig), as shown by its role on patterning of imaginal disc-derived structures in *T. castaneum* [12].

Unhatched embryos from oothecae of dsZelda-treated females showed a wide diversity of malformations. The most frequent were related to abdomen development, followed by morphologically normal first nymphal instar but unable to hatch (Fig 2). When BgZelda-depleted embryos were studied on ED4, a significant number of them showed the development interrupted in very early stages of development, around blastoderm stage, or in earlier stages, around segmentation. A few percentage of embryos were formed in the dorsal part of the egg instead the ventral part. A defect common to all these abnormal embryos was the incompletely formed abdomen, showing the most caudal part amorphous, without traces of segmentation (Fig 3). The importance of Zelda for the development of the posterior zone and the development of the abdomen has been recently described in other short germ-band insects, like the beetle *T. castaneum* and the bug *Rhodnius prolixus* [12].

Our expression studies in BgZelda-depleted embryos suggest that BgZelda promotes the expression of most of the early genes that we examined. The stimulation of the expression of Dnmt1 (Fig 4B) is interesting, as this points to a role of BgZelda in DNA methylation in *B. germanica*. *Dnmt1* is the only DNA methyltransferase gene found in *B. germanica*, and its expression pattern is similar to that of BgZelda, with maximal levels between ED0 and ED2 of embryogenesis and a peak on ED1 [16]. The expression of MIR-309 miRNAs also depends on BgZelda (Fig 4C), a function that is conserved in *D. melanogaster* [2,22]. In *B. germanica*, MIR-309 miRNAs peak on ED2 of embryogenesis [23], that is, one day after the peak of BgZelda (Fig 1F), and we have postulated a role of these miRNAs in eliminating maternal mRNAs during the MZT, as occurs in *D. melanogaster* [32]. Then, all early embryo genes examined (gap, pair-rule and segment polarity genes), were or tended to be down-regulated in BgZelda-depleted embryos (Fig 4D). Previous reports had shown that the expression of most of these genes are affected in DmZelda-deficient *D. melanogaster* [2,6], which suggests that the role of Zelda as a key activator of early zygotic genes [2,6,11] is also present in the less-modified *B. germanica*. We considered that the wrong positioning of the embryo in the dorsal part of the egg may have to do with the expression of genes regulating DV patterning, but none of genes examined was affected in BgZelda-depleted embryos, except *sna* that was significantly down-regulated (Fig 4E). Nevertheless, *sna* is not only involved in the DV patterning [2,29], but it is also required for coordinating mesoderm invagination during gastrulation [33,34]. Our results indicate the BgZelda promotes the expression of *sna*, but we presume that *sna* is not related to the reverse positioning of the embryo but rather to the phenotypes that do not progress beyond blastoderm formation, given its important role in gastrulation.

From an evolutionary point of view, the data suggest that Zelda played the role of activator of the early zygotic genome by binding TAGteam heptamers, in the strict frame of the MZT, in the last common ancestor of modern insects (Neoptera), some 380 million years ago [35]. The occurrence of the canonical TAGteam heptamer CAGGTAG in the genome of Archaeognatha, Odonata and Ephemeroptera (S1 Table), suggests that this role might be already present in the last common ancestor of Insecta (Ectognatha), some 450 million years ago [35]. In any case, the role of zygotic genome activator has been conserved in all major Neopteran clades: Polyneoptera, Paraneoptera and Endopterygota. The contribution to the formation of the abdomen would be also a Zelda function in ancestral Neopterans, which is still present in short germ-band insects, even belonging to Endopterygota. Finally, the expansion of Zelda expression to mid and late embryogenesis that we observe in *D. melanogaster* and *T. castaneum*, with respect to *B. germanica*, might have been instrumental in the innovation of holometaboly in Endopterygota, from hemimetabolan ancestors. We hypothesize that expression of Zelda beyond the MZT in embryo development, would have allowed building the morphologically divergent holometabolan larva, which has been a key step in the evolution of insect metamorphosis from hemimetaboly to holometaboly.

## Materials and Methods

### Insects

Adult females of *B. germanica* were obtained from a colony fed on Panlab dog chow and water *ad libitum*, and reared in the dark at 29 ± 1^°^C and 60–70% relative humidity. Freshly ecdysed adult females were selected and used at appropriate ages. Mated females were used in all experiments, and the presence of spermatozoa in the spermatheca was assessed in all experiments. Prior to injection treatments, dissections and tissue sampling, the animals were anesthetized with carbon dioxide.

### RNA extraction and retrotranscription to cDNA

We performed a total RNA extraction from specific oothecae using the RNeasy Plant minikit (QIAGEN) in the case of early oothecae (since non-fertilized egg until 4 days after ootheca formation, AOF) and GenElute Mammalian Total RNA Miniprep kit in the case of later oothecae (since 6 days AOF to 16 days AOF). In both cases, all the volume extracted was lyophilized in the freeze-dryer FISHER-ALPHA 1–2 LDplus and then resuspended in 8 µl of miliQ H_2_O. For mRNA and miRNA precursors quantification these 8 µl were treated with DNase I (Promega) and reverse transcribed with first Strand cDNA Synthesis Kit (Roche) and random hexamers primers (Roche).

### Quantification of mRNA levels by qRT-PCR

Quantitative real-time PCR (qRT-PCR) was carried out in an iQ5 Real-Time PCR Detection System (Bio-Lab Laboratories), using SYBR^®^Green (iTaq ^TM^Universal SYBR^®^ Green Supermix; Applied Biosystems). Reactions were triplicate, and a template-free control was included in all batches. Primers used to detect mRNA levels studied here are detailed in S4 Table. We have validated the efficiency of each set of primers by constructing a standard curve through three serial dilutions. In all cases, levels of mRNA were calculated relative to BgActin-5c mRNA (accession number AJ862721). Results are given as copies of mRNA of interest per 1000 or per 100 copies of BgActin-5c mRNA.

### RNA interference

The detailed procedures for RNAi assays have been described previously [36]. Primers used to prepare BgZelda dsRNA are described in S4 Table. The sequence was amplified by PCR and then cloned into a pST-Blue-1 vector. A 307 bp sequence from *Autographa californica* nucleopolyhedrosis virus (Accession number K01149.1) was used as control dsRNA (dsMock); primers used to synthetize dsRNA are also described in S4 Table. The dsRNAs were prepared as reported elsewhere [36]. A volume of 1 µl of the dsRNA solution (3 µg/µl) was injected into the abdomen of 1-day-old adult females, with a 5µl Hamilton microsyringe. Control specimens were treated at the same age with the same dose and volume of dsMock.

### Microscopy

Oothecae were detached from the female abdomen by gentle pressure or obtained directly because the animal left them. Each ootheca was opened and the embryos were dechorionated and individualized. Then these embryos were directly observed under the stereo microscope Carl Zeiss – AXIO IMAGER.Z1. For DAPI staining, after 10 min in a water bath at 95ºC, each ootheca was opened and the embryos dechorionated and individualized. Between 12 and 24 embryos per ootheca, chosen from the central part, were dissected for staining. These embryos were fixed in 4% paraformaldehyde in PBS for 1h, then washed with PBS 0.3% Triton (PBT) and incubated for 10 min in 1 mg/ml DAPI in PBT. They were mounted in Mowiol (Calbiochem, Madison, WL, USA) and observed with the fluorescence microscope Carl Zeiss – AXIO IMAGER.Z1.

### Statistical analyses of qRT-PCR

In all experiments, to test the statistical significance between treated and control samples it has been used the Relative Expression Software Tool (REST), which evaluates the significance of the derived results by Pair-wise Fixed Reallocation Randomization Test [37].

### Transcriptomic and genomic data

We obtained the transcriptome-based pattern of expression of Zelda in *B. germanica, D. melanogaster* and *T. castaneum*. Those of *B. germanica* and *D. melanogaster* were identical to those previously obtained by Ylla et al. [16], who precisely describe the different stage-libraries used. The RNAseq dataset of *B. germanica* and *D. melanogaster* are are accessible at GEO: GSE99785 [16] and GEO: GSE18068 (Celniker et al., 2009; modENCODE Consortium et al., 2010), respectively. The RNA-seq dataset of *T. castaneum* embryogenesis used (GEO: GSE63770) comprises 8 libraries from 4 developmental stages (2 replicates each) covering non-fertilized eggs (NFE), and three sequential embryo stages: 8–16 h, 16–24h, 24–48h [19]. In addition, we obtained a RNA-seq library from *T. castaneum* adult females [38] available at SRA: SRX021963. The TAGteam heptamers were examined along the genome assemblies and its complementary sequences using custom-made Python scripts. The calculation of heptamer densities within scaffolds and correlations between genome length and number of heptamers was preformed within R. The complete list of genomes used and their accession can be found in the S1 Table.

## Supporting information

**S1 Fig.** Alignment of the C-terminal region, containing the four C2H2 zinc fingers (ZF3 to ZF6) of the Zelda protein sequence of *Blattella germanica* (BgZelda) and *Drosophila melanogaster (DmZelda)*. The percentage of similarity in this region is 76.7%.

**S2 Fig.** Number of CAGGTAG motifs within 10 Kb windows along each scaffold of the genome sequence of *Blattella germanica*. The abscissae show the scaffold position, and the ordinate shows the number of motifs in the given 10 Kb window. Only the first 15 scaffolds are represented, but the tendency is similar in all scaffolds. The scaffold number refer to the genome version available at NCBI Bioproject, accession code PRJNA203136.

**S3 Fig.** Transcriptomic expression of *zelda* in *Blattella germanica, Drosophila melanogaster* and *Tribolium castaneum*. Data of *B. germanica* and *D. melanogaster* are as in Ylla et al. (2018). In *B. germanica*, the following 11 stages are represented: non-fertilized egg (NFE), 8, 24, 48, 144 and 312 hours after oviposition (Embryo 0, Embryo 1, Embryo 2, Embryo 6 and Embryo 13), first, third, fifth and sixth (last) nymphal instars (Nymph 1, Nymph 3, Nymph 5 and Nymph 6) and the Adult (female) (Ylla et al., 2018). In *D. melanogaster, the following 11 stages are represented: six* sequential embryo stages (Embryo 0–4h, Embryo 4–6h, Embryo 6–12h, Embryo 12–16h, Embryo 16–20h, Embryo 20–24h), the three larval stages (Larva 1, Larva 2, Larva 3), the Pupa, and the Adult (female) (Celniker et al., 2009; modENCODE Consortium et al., 2010). In *T. castaneum*, the following 5 stages are represented: non-fertilized eggs (NFE) and three sequential embryo stages: 8–16h, 16–24h, 24–48h (Ninova et al., 2016), and the Adult (female) (Altincicek et al., 2013).

**S1 Table.** Insect genomes used for correlating the number of canonical Zelda-binding heptamer CAGGTAG and the genome size.

**S2 Table.** Scaffolds of the *Blattella germanica* genome and CAGGTAG motifs in them.

**S3 Table.** Number of TAGteam motifs in the genes examined in Zelda-depleted embryos. We considered the number of motifs present inside the gene sequence. The scaffold number refer to the *Blattella germanica reference*.

**S4 Table.** Primers used to to measure expression levels by qRT-PCR. In the case of *zld*, we also indicate the primers used to prepare the dsRNA for RNAi experiments. *Genes manually annotated in *Blattella germanica* genome, available as BioProject PRJNA203136.

## Acknowledgements

This work was supported by the Spanish Ministry of Economy and Competitiveness (grants CGL2012–36251 and CGL2015–64727-P to X.B. and predoctoral fellowship to A.V.-A.) and the Catalan Government (grant 2017 SGR 1030 to X.B.). It also received financial assistance from the European Fund for Economic and Regional Development (FEDER funds). Natalia Llonga helped to find TAGteam motifs associated to the genes examined in the expression studies.

